# Analyzing SARS CoV-2 Patient Data Using Quantum Supervised Machine Learning

**DOI:** 10.1101/2021.10.26.466019

**Authors:** Zara Yu

**Affiliations:** Department of Physics, Stevens Institute of Technology, Castle Point on the Hudson, Hoboken, NJ 07030 and Kinnelon High School, Kinnelon, NJ 07405

## Abstract

The novel coronavirus disease 2019 (COVID-19) has created a serious threat to global health. We developed a new quantum machine learning (QML) assisted diagnostic method that can provide an accurate diagnosis to aid decision processes of medical providers. One of the key elements in our method was to implement the quantum variational method to efficiently classify data, taking crucial multiple correlations among the features into account. We established and fine-tuned this quantum classifier by using a group of data drawn from publicly available COVID-19 cases. We have shown that QML is capable of processing patient information efficiently and accurately for the diagnosis of COVID-19.

## I. Introduction

The virus SARS-CoV-2 (commonly known as COVID-19) has led to a worldwide pandemic which has jeopardized the health of millions globally. The virus first appeared in early December of 2019 in Wuhan, China where it quickly spread to countries around the world. From August 2021, the total number of patients that have been confirmed to have COVID-19 is over 206,000,000 and the disease has affected over 200 countries. Over 4,000,000 people have died because of the disease. The pandemic has disrupted healthcare systems all over the world. For example, new plans had to be developed to accommodate the decrease in number for available acute care beds and supply of personal protective equipment (PPE) had to be monitored carefully. During this time, healthcare workers needed to learn how to make quick clinical decisions and use resources productively.

SARS-CoV-2 causes those who are affected to exhibit a large variety of symptoms including fever, cough, chest pain, shortness of breath, and gastrointestinal issues among many others. The point of care test is highly important for diagnosis, but the testing process is time-consuming, and not always reliable. It has been reported in many cases that false positive or false negative results are not uncommon, particularly in the early stages of the illness. For a contagion with rapid transmission like COVID-19, it is difficult to discern if a patient in the early stages has contracted SARS-CoV-2 or if the symptoms are caused by a different virus. Therefore, developing an accurate and efficient diagnostic system is highly desirable when a scientifically sound decision is needed to triage patients and provide the appropriate medical care.

Artificial intelligence (AI), expert systems (ES) and machine learning (ML) algorithms have been used in medical diagnosis methods for a long time. For example, ML and AI have been used to improve the accuracy and speed of several screening processes, while dealing with big data issues. But for a worldwide pandemic like SARS-CoV-2, new challenges are posed such as the surge in suspected cases, the lack of reliable data, inconclusive diagnoses, and the necessity for rapid diagnosis. Quantum physics is known to be useful in speeding up the information processing rate. The application of quantum theory, particularly quantum coherence and entanglement to machine learning algorithms, has substantially improved speed and accuracy in other arenas. Therefore, we would like to test this in the area of medical diagnosis. We first theoretically formulated a mathematical model that naturally incorporated both conventional machine learning and the quantum inspired components. The method used in this project combines three different important elements to give rise to a type of diagnostic method based on a quantum machine learning method. We proposed, following a new type of quantum machine learning, to use a quantum variational method to generate a new type of identification procedure that can be particularly useful for an early and rapid diagnosis. This was achieved by employing quantum gates in an n-qubit system. Thus, the quantum variational method may be realized as the combination of traditional pattern recognition processes and quantum operations. In our project, the data was coded by the standard states of a quantum mechanical system represented by multi-dimensional vectors. The quantum states carrying the coded information are manipulated according to quantum mechanics rules to maximize the efficiencies. For example, quantum inspired methods allowed us to consider not only multiple symptoms simultaneously such as temperature, fatigue, cough, shortness of breath, muscle aches, headache, loss of taste or smell, nausea etc., it also included the correlation among those symptoms. This is only an example of a much general method, in which all the provided symptoms and associated diagnosis criteria will be analyzed systematically through fast numerical algorithms combined with certain well-defined quantum operations and measurements.

The purpose of this research project is to develop a new quantum ML assisted diagnostic method that can process crucial information efficiently and provide a rapid diagnosis to aid medical providers’ decision processes. We want to emphasize that the mathematical model used in this paper is not designed to be used as a substitute for standardized diagnostic methods for COVID-19 including the NAAT (nucleic acid amplification test) and antigen tests. Furthermore, it does not replace an appropriate physical examination or critical analysis in a clinical setting. Our main goal of this research is to provide a useful diagnostic tool to assist medical providers in identifying patients during the triage process who may need priority care, including a COVID-19 test. This can be particularly important when the sensitivity of the test is low or in cases when medical resources and hospital capacity are limited.

The paper is organized as follows. We introduce our methodology and data in Sec. II. The numerical simulations and results are given in Sec. III. In this section, we also investigate how to interpret our results in their applications to COVID-19 diagnosis. We comment and conclude in Sec. IV. The appendix gives a simple example of a quantum variational classifier and describes some basic ideas of the classifier.

## II. Methodology

The so-called quantum learning method used in this paper is based on PennyLane which is an efficient program of implementing variational quantum classifiers [1–3]. The essence of variational quantum classifiers is to train quantum circuits by using a set of known data samples to classify new data sets. Please see the appendix for an example of a quantum variational classifier.

In this paper, we show how to train the COVID-19 model to recognize the two classes of data for four different features. We have reformulated the quantum variational method to make it applicable for SARS-CoV-2 problems. We included the correlation information among different symptoms. Then, we found an efficient way to simulate these correlation functions. Using the Pennylane program (freely available on Pennylane.ai) and the Anaconda Python application, we were able to design a properly working quantum variational classifier. The classifier was used to groups data into multiple features. Here, we only considered two major features, but a more detailed investigation with more features will be reported in the future. Initially, the Iris data set was used to test the classifier and determine if it worked. Before deciding on using the variational classifier, other classifiers such as a data re-uploading classifier and an ensemble classifier were evaluated as well.

In order to test our model, we collected a set of real data that was publicly available. The data was fed into our model to train our model. Typically, the theoretical model will give better performance with the increased data inputs. The data collection was conducted through online searching to find the publicly available information from websites and published research papers that had included real patient data for SARS-CoV-2 and were available to the public. The websites included government (for example CDC), education, and health research institution websites. We follow the research ethical standards to conduct our research. We used datasets from the Influenza Research Database from fludb.org [12], the National Institute of Health, and the Israeli Ministry of Health website [17]. We also tested the model with different sets of data with multiple criteria in mind, including efficiency and accuracy. The methods in this part are to use fidelity in distinguishing quantum states, and we also estimated the complexity associated with quantum operations and quantum measurements.

## III. Results and Discussions

We use the quantum variational classifier to generate multiple graphs that can exhibit the interesting relationship between the studied cases and the chosen features such as temperature (fever), fatigue, muscle pain, coughing etc. As we have mentioned before, the aim of this project is to investigate the correlation between the studied cases and the features.

Figs. 1–3 are used to exhibit the correlation between the chosen features for four sets of data. For simplicity, we only plot three out of four features. For Fig. 1, the x-axis (horizontal) represents the temperature of each data point while the y-axis (vertical) represents the feature or severity of one of the symptoms that was measured (here we highlight runny nose severity). In Fig. 2, we study the correlations between two features involving muscle pain severity and temperature. In these plots, the chosen features include the following symptoms: temperature, runny nose, and muscle pain. Fig. 3 is plotted to show the corrections between the chosen features and the Covid-19 case diagnosis results.

**FIG. 1:**
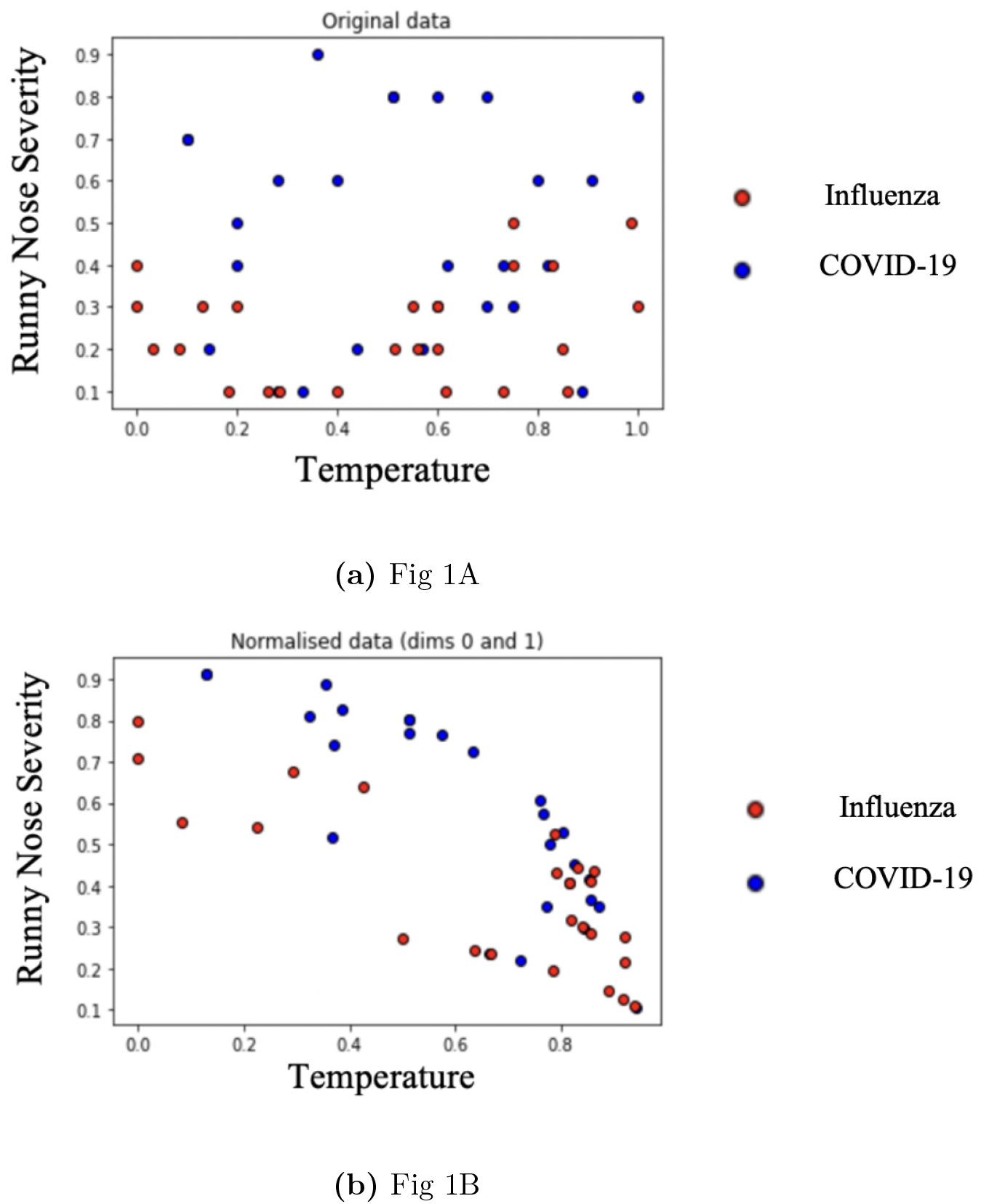
The figures show the two plots initially generated from Dataset 1. In Fig. 1A, the data that has not been normalized is plotted. In Fig. 1B, the data is normalized to be appropriate for the numerical simulations.

**FIG. 2:**
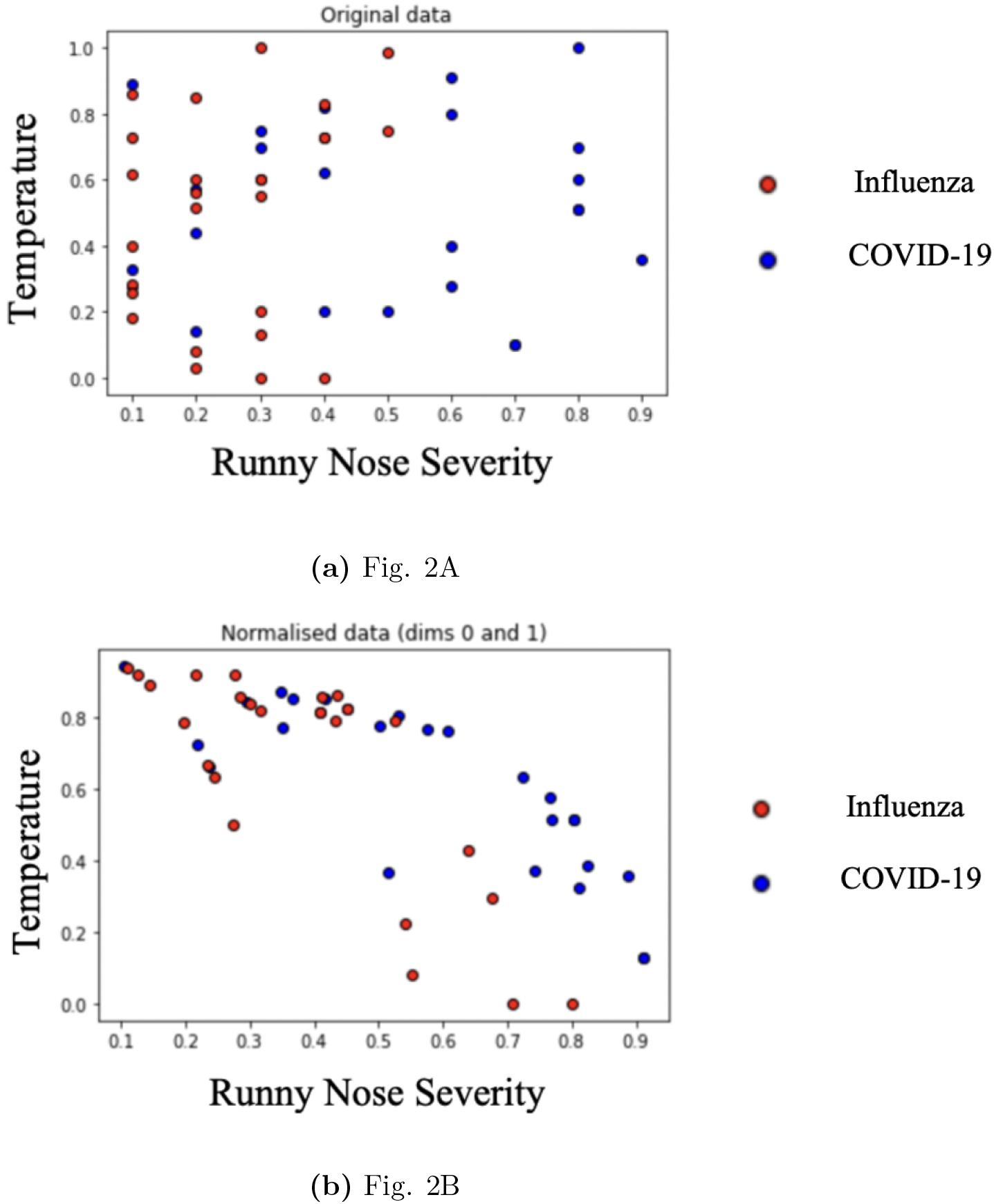
The figures show the plots initially generated from Dataset 2. In Fig. 2A, the data that has not been normalized is plotted. In Fig. 2B, the data is normalized to be appropriate for the numerical simulations.

**FIG. 3:**
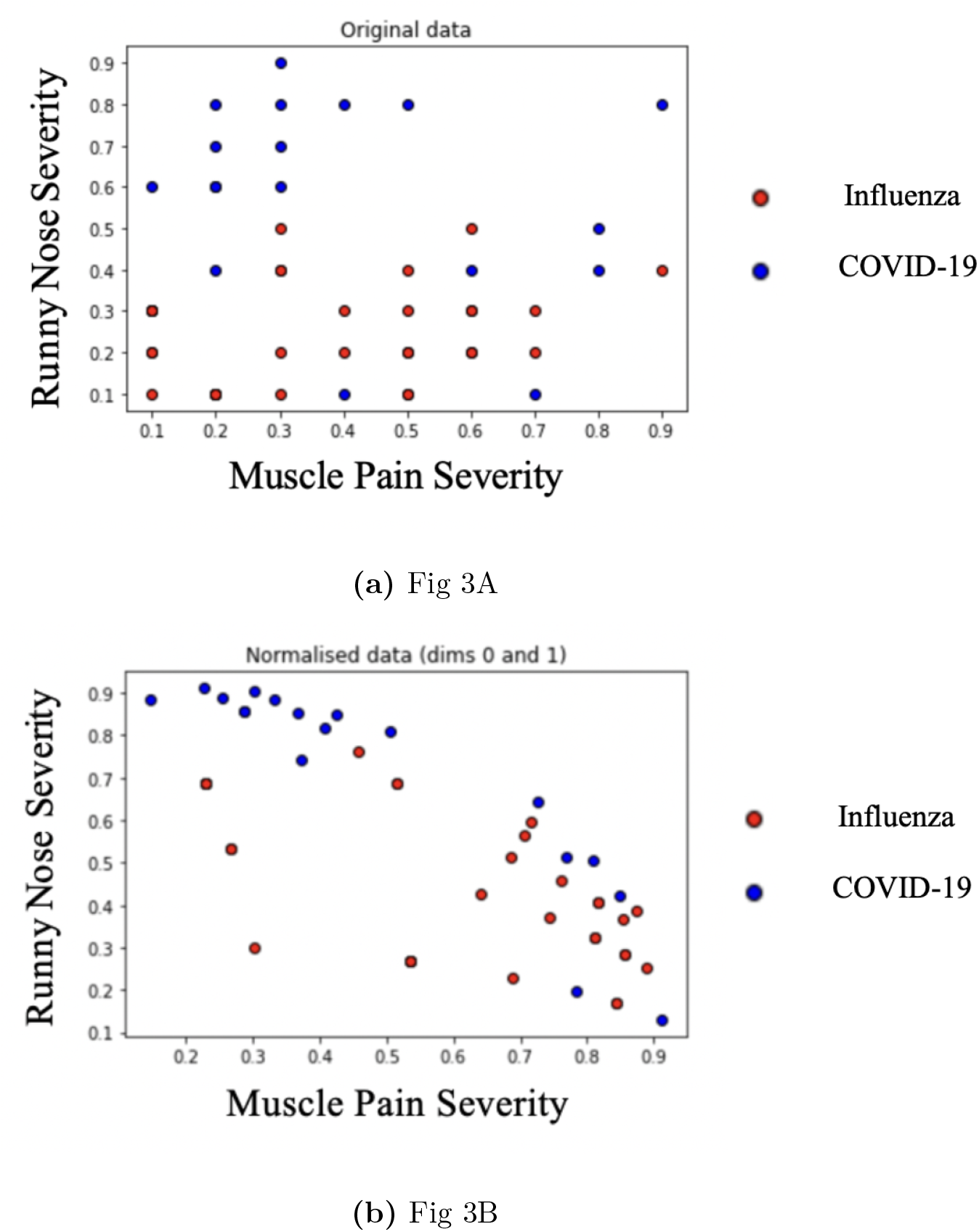
The figures show the plots initially generated from Dataset 3. In Fig. 3A, the data that has not been normalized is plotted. In Fig. 3B, the data is normalized to be appropriate for the numerical simulations.

The temperature range of the data collected is typically from 97 to 103 degrees Fahrenheit. For the numerical simulations, the temperature range is converted into angle degrees from 0 to 180 degrees and then normalized. For example, the temperatures 97, 97.1, 97.2 are assigned numbers 0, 3, 6. Then the numbers assigned are divided by 180 in order to get a normalized data set. As shown above, all the data used for the plots are normalized. It should be noted that some features such as the severity of symptoms cannot be easily represented numerically. However, in order to feed the data to our model system and carry out the numerical simulations efficiently, we have to assign numbers to the chosen features that are not naturally measured by numbers. According to our convention, if the patients were said to have no symptoms, they were assigned a random number from 0-0.3. If the patients had mild symptoms, they were assigned a random number from 0.4-0.7. If they patient had severe symptoms, they were assigned a random number from 0.8-1.0. Of course, this numbering system is certainly not perfect, but it is useful to numerically simulate our model that will process real cases. We believe that a more systematic method to convert the degrees of severity to a set of numbers is certainly desirable. For example, it is easy to see that the model can be significantly improved when the numbers are assigned by the experts and medical providers. In addition, it is clear that, when a large amount of data become available, this subjective number-assigning system will become more objective.

In Figs. 1-3, two figures (A, B) are plotted using our quantum inspired classifier against the two features represented by the x-axis (horizontal) and y-axis (vertical), respectively. The features that are plotted are decided by the ordering of the data when inputted into the code. For example, if the temperature feature was inputted first, it would be plotted along the x-axis. Then, if the severity of muscle pain was inputted next, it would be plotted along the y-axis. The rest of the features would be used in determining the separation on the graph. We see clearly that the model has exhibited several very interesting features. The model based on our quantum inspired classifier can indeed predict some useful correlations between a patient case with SARS-CoV-2 and the chosen features. Of course, the model can be optimized if more data are fed into the system.

In Fig. 4, the original data is once again plotted but the normalized data and feature vectors are taken into account when shading the graph. This creates distinction between the SARS-CoV-2 and influenza sections. The darker red areas of the graph represent spots where patient points are very likely to have influenza. As the color becomes lighter, it means there’s a lower probability. As the graph becomes blue, it indicates that patient data points are more likely to have. For Fig. 4A, the features that are used in the x-axis and y-axis are temperature and runny nose severity, respectively. For Fig. 4B, the features that are used in the x-axis and y-axis are runny nose severity and temperature, respectively. For Fig. 4C, the features that are used in the x-axis and y-axis are muscle pain severity and runny nose severity, respectively.

**FIG. 4:**
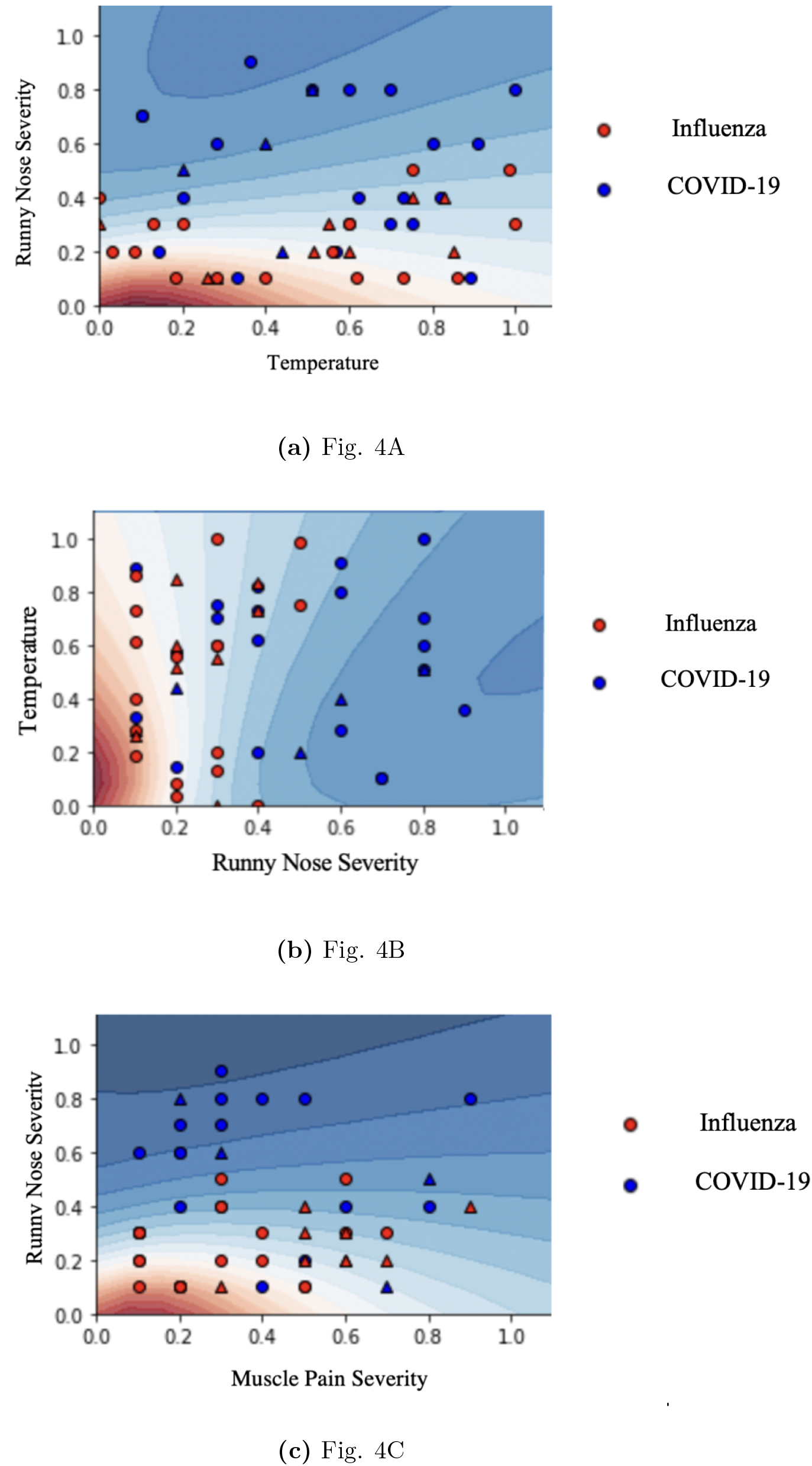
The continuous output of the quantum variational classifier from PennyLane with Datasets 1-3.

As seen from Fig. 4, the model adapted in this paper can be used to separate the confirmed cases from the suspected ones with good to excellent prediction. Based on our model, when data points are closer to the darker red zones, it means that they are more likely to have influenza. On the other hand, when data points are closer to the darker blue zones, this indicates that they are more likely to have SARS-CoV-2 affection. We can see that some data are in regions of opposite color; for such a mathematical model, one cannot expect that the model can make a perfect separation between SARS-CoV-2 and influenza.

As mentioned above, the goal of this project was to show how to use QML to assist COVID-19 pre-testing diagnosis processes. Given our datasets, the quantum inspired machine learning method has been shown to be capable of providing efficient and reasonably accurate results.

## IV. Conclusions and Outlook

We have shown that the quantum inspired ML method can provide an efficient and accurate analysis for the COVID-19 diagnosis processes. High efficiency is clearly desirable for combating the threat of a global crisis like the SARS-CoV-2 pandemic. In such situation, rapid data collection, information processing, and reliable diagnosis processes are essential. The advantage of our method is the implementation of the quantum variational method that has proven to be efficient in classifying data, taking account of crucial multiple correlations. With our chosen features, we have shown that quantum inspired machine learning can be efficient and accurate in information processing and diagnosing COVID-19 cases. Future work consists of fine-tuning our quantum inspired model with more data inputs and multiple chosen features. Ultimately, the goal is to implement the proposed method in clinical diagnosis by developing a practical software package.

In addition to quantum ML methods, we may explore how to incorporate quantum correlation such as quantum entanglement to accelerate the simulation processes. With increased numbers of data, we expect the model can be tuned to provide more reliable predictions. Particularly, we will use a quantum trajectory approach to simulate multi-dimensional systems. The advantage of the quantum trajectory approach is that the computer can efficiently deal with many small “tedious” tasks to give rise to a satisfactory solution through expectation values.

## V. Acknowledgements

We thank Dr. Rupak Chatterjee for his useful discussions on quantum machine learning.

## VI. Appendix

Let us consider a quantum computation process involving a set of unitary matrices

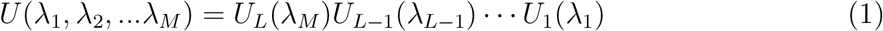

where the control parameters are represented by λ_1_,λ_2_,…λ_*M*_. For simplicity, we only use one parameter to define individual unitary matrices. We may use |*ψ*〉 to denote the initial state labeled by a set of input data,

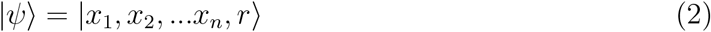

where the data set *x* = *x_1_x_2_*…*x_n_*, and each *x_i_* takes the value +1 or 1, the qubit |*r*〉 is a readout qubit. The final state after a sequence of quantum operations |*ψ*〉_*f*_ will be given by

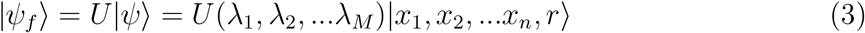

The basic ideas underlying the variational quantum classifiers is to find an optimal parameter set for λ_*i*_ by feeding the known data to the system and measuring the outputs. The training processes are implemented by essentially converting a classification problem into an optimization problem through introducing cost functions that might be defined as a measure of the distance between the target labels and the model outputs. The loss function may be defined as follow [1],

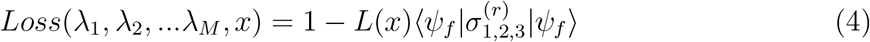

where 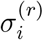 (*i* = 1,2,3) are Pauli matrices acting on the read-out qubit |*r*〉 and the binary label *L*(*x*) is chosen as +1 or 1. Physically, 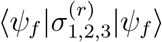 represents the mean values of measuring Pauli matrices on the read-out state.

